# Ultrafast Simulation of Large-Scale Neocortical Microcircuitry with Biophysically Realistic Neurons

**DOI:** 10.1101/2021.02.22.432356

**Authors:** Viktor János Oláh, Nigel P Pedersen, Matthew JM Rowan

## Abstract

Understanding the activity of the mammalian brain requires an integrative knowledge of circuits at distinct scales, ranging from ion channel gating to circuit connectomics. To understand how multiple parameters contribute synergistically to circuit behavior, neuronal computational models are regularly employed. However, traditional models containing anatomically and biophysically realistic neurons are computationally demanding even when scaled to model local circuits. To overcome this limitation, we trained several artificial neural net (ANN) architectures to model the activity of realistic, multicompartmental neurons. We identified a single ANN that accurately predicted both subthreshold and action potential firing and correctly generalized its responses to previously unobserved synaptic input. When scaled, processing times were orders of magnitude faster compared with traditional approaches, allowing for rapid parameter-space mapping in a circuit model of Rett syndrome. Thus, we present a novel ANN approach that allows for rapid, detailed network experiments using inexpensive, readily available computational resources.

## Introduction

Understanding the behavior of complex neural circuits like the human brain is one of the fundamental challenges of this century. Predicting mammalian circuit behavior is difficult due to several underlying mechanisms at distinct organizational levels, ranging from molecular-level interactions to large-scale connectomics. Computational modeling has become a cornerstone technique for deriving and testing new hypotheses about brain organization and function^1-4^. In little more than 60 years, our mechanistic understanding of neural function has evolved from describing action potential (AP) related ion channel gating^5^ to constructing models that can simulate the activity of whole brain regions^6-10^. Although tremendous advancements have been made in the development of computational resources, the lack of available or affordable hardware for neural simulations currently represents a significant barrier to entry for most neuroscientists and renders many questions intractable. This is particularly well illustrated by large-scale neural circuit simulations. In contrast to detailed single-cell models, which have been a regular occurrence in publications since the ‘90s^11-18^, parallel simulation of thousands, or even hundreds of thousands of detailed neurons have only become a possibility with the advent of supercomputers^19-26^. As these resources are still not widely accessible, several attempts have been made to mitigate the immense computational load of large-scale neural simulations by judicious simplification^19,27-33^. However, simplification inevitably results in feature or information loss, such as sacrificing multicompartmental information for simulation speed^19,27,28,30^. Thus, there is a critical need for new approaches to enable efficient large-scale neural circuit simulations on widely available computational resources without surrendering biologically relevant information.

To counteract the increasing computational burden of ever-growing datasets on more traditional models, many fields have recently adopted various machine learning algorithms^34-38^. Specifically, artificial neural networks (ANNs) are superior to conventional model systems both in terms of speed and accuracy when dealing with complex systems, such as those governing global financial markets or weather patterns^39,40^. Due to their accelerated processing speed, ANNs are ideal candidates for modeling large-scale biological systems. The idea that individual neural cells could be represented by ANNs was proposed almost two decades ago^41^, however, current ANN solutions are still unfit to replace traditional modeling systems as they cannot generate gradational neuronal dynamics needed for network simulations. Therefore, we aimed to develop an ANN that can (1) accurately replicate various features of biophysically detailed neuron models, (2) efficiently generalize for previously unobserved input conditions and (3) significantly accelerate large-scale network simulations.

Here we investigated the ability of several ANN architectures to represent membrane potential dynamics, in both simplified point neurons and multicompartment neurons. Among the selected ANNs, we found that a convolutional recurrent architecture can accurately simulate both subthreshold and suprathreshold voltage dynamics. Furthermore, this ANN could generalize to a wide range of input conditions and reproduce neuronal features following different input patterns beyond membrane potential responses, such as ionic current waveforms. Next, we demonstrated that this ANN could also accurately predict multicompartmental information, by fitting this architecture to a biophysically detailed layer 5 pyramidal cell^42^ model, as well as several other distinct cortical neurons. Importantly, we found that ANN representations could exponentially accelerate large network simulations, as demonstrated by network parameter space mapping of a cortical recurrent microcircuit example model of Rett syndrome, a neurodegenerative disorder associated with cortical dysfunction and seizures^43-46^. Thus, we provide a detailed description of an ANN architecture suitable for large-scale simulations of anatomically and biophysically complex neurons, applicable to human disease modeling. Most importantly, our ANN simulations are accelerated to the point where detailed network experiments can now be carried out using inexpensive, readily available computational resources.

## Results

To create a deep learning platform capable of accurately representing the full dynamic membrane potential range of neuronal cells, we focused on model systems proven to be suitable for multivariate time series forecasting (MTSF). To compare the ability of different ANNs to reproduce the activity of an excitable cell, we designed five distinct architectures (Fig. 1a). The first two models were a simple linear model with one hidden layer (linear model, Fig. 1a, blue) and a similar model equipped with nonlinear processing (nonlinear model, Fig. 1a, cyan), as even relatively simple model architectures can explain the majority of subthreshold membrane potential variance^47^. The third and fourth models consist of recently constructed time-series forecasting architectures, including a recurrent ANN (CNN-LSTM, Fig. 1a, magenta) consisting of convolutional layers^48^, long short-term memory (LSTM^49,50^) layers, and fully connected layers, termed the CNN-LSTM network (Supplementary Fig. 1,^51^) and a more recently developed architecture relying on dilated temporal convolutions (convolutional net, Fig. 1a, orange) (based on the WaveNet architecture^52,53^), which is superior to the CNN-LSTM in several MTSF tasks. The CNN-LSTM has the distinct advantage of having almost two orders of magnitude more adjustable parameters compared to the aforementioned ANNs. Finally, we selected a fifth architecture (deep neural net, Fig. 1a, green) with a comparable number of free parameters to the CNN-LSTM, composed of ten hidden layers, which operates solely on linear and nonlinear transformations. Before moving to neural cell data, each of the five selected architectures were evaluated using a well-curated weather time series dataset (see methods). Each model performed similarly (0.070/0.069, 0.059/0.06, 0.089/0.094, 0.07/0.069, 0.092/0.095, mean absolute error on the validation/testing datasets for linear, nonlinear, convolutional net and CNN-LSTM, deep neural net architectures, respectively), demonstrating their suitability for MTSF problems.

**Fig. 1.**
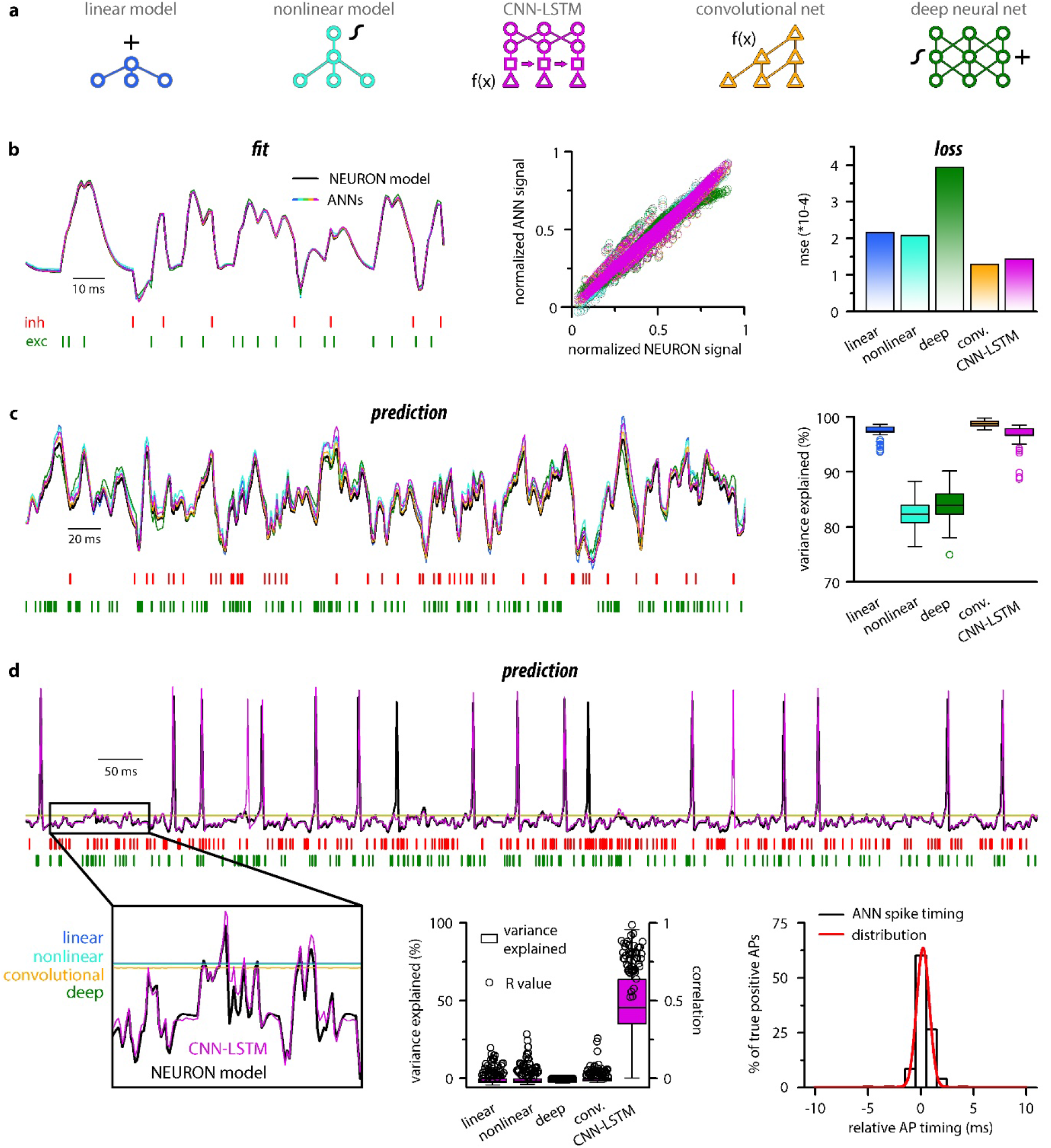
Single compartmental neuronal simulations using ANNs. **a**. Representative diagrams of the tested architectures, outlining the ordering of the specific functional blocks of the ANNs. **b**. Point-by-point fit of passive membrane potential by ANNs as a continous representative trace (left) and plotted agains ground truth data (middle, n = 45000). Mean squared error of ANN fits corresponds to the entire training dataset (n = 2.64*10^6^ datapoints). Single quantal inputs arrive stochastically with a fixed quantal size: 2.5 nS for excitatory, 8 nS for inhibitory inputs, sampling is 1 kHz. Red and green bars below membrane potential traces denote the arrival of inhibitory and excitatory events, respectively. **c**. Representative trace of a continous passive membrane potential prediction (left) created by relying on past model predictions. Explained variance (right) was calculated from 500 ms long continous predictions (n = 50). **d**. Representative active membrane potential prediction by ANNs (top, left). Explained variance (middle, box chart) and Pearson’s r (middle, circles) of model predictions and ground truth data for the five ANNs from 50 continous predictions, 500 ms long each. Spike timing of the CNN-LSTM model calculated from the same dataset as middle panel. Color coding is the same as in panel **a**.

### Prediction of point neuron membrane potential dynamics by ANNs

To test the ability of the five ANNs to represent input-output transformations of a neural cell, we next fitted these architectures with data from passive responses of a single-compartmental point-neuron model (NEURON simulation environment^54^) using the standard backpropagation learning algorithm for ANNs^55^. Each model was tasked with predicting a single membrane potential value based on 64 ms (a time window that yielded the best results both in terms of speed and accuracy) of preceding membrane potentials and synaptic inputs (Fig. 1). We found that both the linear and nonlinear models predicted subsequent membrane potential values with low error rates (Fig. 1B) with similar behavior in both the CNN-LSTM and convolutional architectures (2.16*10^−4^ ± 1.18*10^−3^, 2.07*10^−4^ ± 1.11*10^−3^, 1.43*10^−4^ ± 9.31*10^−4^, 1.29*10^−4^ ± 9.42*10^−4^ mean error for linear, nonlinear, CNN-LSTM and convolutional models, respectively). However, the deep neural network performed considerably worse than all other tested models (3.94*10^−4^ ± 1.56*10^−3^ mean error), potentially due to the nonlinear correspondence of its predicted values to the ground truth data (Fig. 1b).

Next, we tested ANNs in simulation conditions similar to the traditional models. To this end, we initialized ANNs with ground truth data followed by a continuous query period in which forecasted membrane potential values were fed back to the ANNs to observe continuous unconstrained predictions. As expected from the fit error rates of single membrane potential forecasting (Fig. 1b), continuous predictions of the linear, convolutional, and CNN-LSTM models could explain the ground truth signal variance at high accuracy. At the same time, the deep neural net performed slightly worse (Fig. 1c, 0.97 ± 0.01%, 0.99 ± 0.01%, 0.97 ± 0.02%, 0.84 ± 0.03% variance explained for linear, convolutional, CNN-LSTM, and deep neural net architectures respectively, n = 50). Surprisingly, the nonlinear model produced the worst prediction for passive membrane potential traces (0.82 ± 0.03% variance explained, n = 50) despite performing the best on the benchmark dataset. Together, these results indicate that even simple linear ANNs can capture subthreshold membrane potential behavior accurately^47^.

Next, we tested how these models perform on the full dynamic range of neural cells, which due to AP firing (which can also be viewed as highly relevant outlier datapoints), constitutes a non-normally distributed and thus demanding dataset for ANNs. Interestingly, we found that only the CNN-LSTM architecture could precisely reproduce both subthreshold membrane potential dynamics and spiking activity, while all other tested ANNs converged to the mean of the training dataset (Fig. 1d, -0.027 ± 0.073%, -0.025 ± 0.068%, -0.018 ± 0.054%, 0.456 ± 0.304%, -0.025 ± 0.067 % variance explained for linear, nonlinear, convolutional net and CNN-LSTM, deep neural net architectures respectively, n = 50). We found that although the CNN-LSTM model explained substantially less variance for the active membrane potential traces (Fig. 1d) than for subthreshold voltages alone (Fig. 1c), the predictions showed high linear correlation with the ground truth signals (Pearson’s r = 0.76793 ± 0.10003, n = 50). For the four remaining ANN architectures, it is unlikely that convergence to the mean is caused by settling in local minima on the fitting error surface as ANNs have a large number of free parameters (2.07*10^4^, 2.07*10^4^, 2.47*10^6^, 3.64*10^5^, 1.95*10^6^ free parameters for linear, nonlinear, deep, convolutional ANNs and CNN-LSTM respectively). Therefore the chance of having a zero derivative for each parameter at the same point is extremely low^56^, suggesting that erroneous fitting is the consequence of the limitations of these ANN architectures. Consequently, of the tested ANN architectures, the CNN-LSTM is the only model that could depict the full dynamic range of a biophysical neural model.

**Supplementary Fig. 1.**
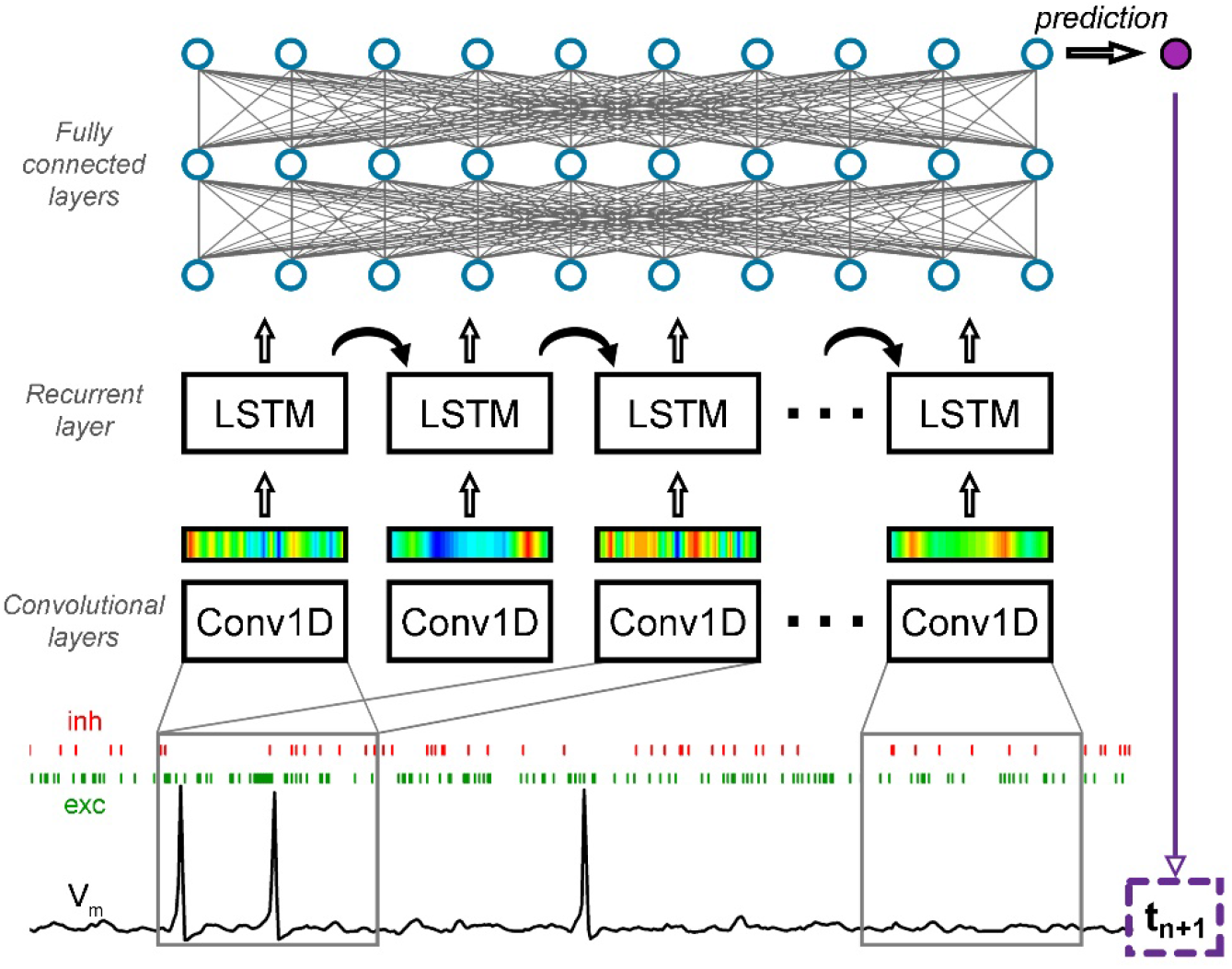
CNN-LSTM architecture for time series forecasting. The input of the ANN consisted of a membrane potential vector (V_m_) and the weights and onsets of synaptic inputs (a representative inhibitory synapse – inh in red, and an excitatory synapse – exc in green). The first layer of the ANN (Conv1D) creates a temporally alligned convolved representation (colored bars) of the input by sliding a colvolutional kernel (grey box) along the input. The second functional block (LSTM layers) processes the output of the convolutional layers through recurrent connections, to weigh information temporally. The last functional block consisting of fully connected layers provides additional nonlinear information processing power. The output of the network in this case is the first subsequent V_m_ value (t_n+1_). The number of layers belonging to specific functional blocks of the CNN-LSTM architecture may vary. Red and green bars below membrane potential traces denote the arrival of inhibitory and excitatory events, respectively.

Closer inspection of the timing of the predicted APs revealed that the CNN-LSTM models correctly learned thresholding, as the occurrence of the APs matched the timing of the testing dataset (Fig. 1d; 83.94 ± 16.89% precision and 90.94 ± 12.13 % recall, 0.24 ± 0.79 ms temporal shift for true positive spikes compared to ground truth, n = 283). Taken together, we developed an ANN architecture that is ideally suited for predicting both subthreshold membrane potential fluctuations and the precise timing of APs on a millisecond timescale.

### Generalization of the CNN-LSTM architecture

To test the applicability of the CNN-LSTM for predicting physiological cellular behavior, we assessed the generalization capability of the architecture built for active behavior prediction (Fig. 1d). Generalization is the ability of an ANN to accurately respond to novel data^57,58^. According to our hypothesis, if the CNN-LSTM correctly learned the mechanistic operations of a neural cell, then the architecture should behave appropriately when tasked with responding to novel quantal amplitudes and input patterns.

We first challenged the CNN-LSTM by administering excitatory inputs with variable quantal sizes (0.1-3.5 nS, 0.1 nS increment). Similar to the control NEURON model, the CNN-LSTM responded linearly in subthreshold voltage regimes (Fig. 2a, Pearson’s r = 0.99, n=35) and elicited an AP after reaching threshold. Independent evaluation of the NEURON model control revealed a surprisingly similar I/V relationship for the same quantal inputs (intercept, -0.003 ± 8.53 and -0.003 ± 0.001; slope for subthreshold linear I/V, 22.2 ± 0.41 and 23.31 ± 0.62; CNN-LSTM and NEURON model, respectively) and similar AP threshold (−58.03 mV and -56.64 mV for CNN-LSTM and NEURON model, respectively). Next, we tested temporal summation of excitatory inputs (Fig. 2b). We found that the independently simulated NEURON model displayed similar temporal summation patterns to the CNN-LSTM both for sub- and suprathreshold events (Fig. 2b). Finally, we combined the previous two tests and delivered unique temporal patterns of synaptic inputs with variable synaptic conductances randomly chosen from a normal distribution (mean: 2.5 nS, variance: 0.001 nS, Fig. 2c). Again, the predictions of the CNN-LSTM architecture closely matched traces obtained from the NEURON model (Pearson’s r = 0.81, n = 5000 ms) and the timing of the majority of the APs agreed with the ground truth data (91.02 ± 16.03 % recall and 69.38 ± 22.43 % precision, n = 50).

**Fig. 2.**
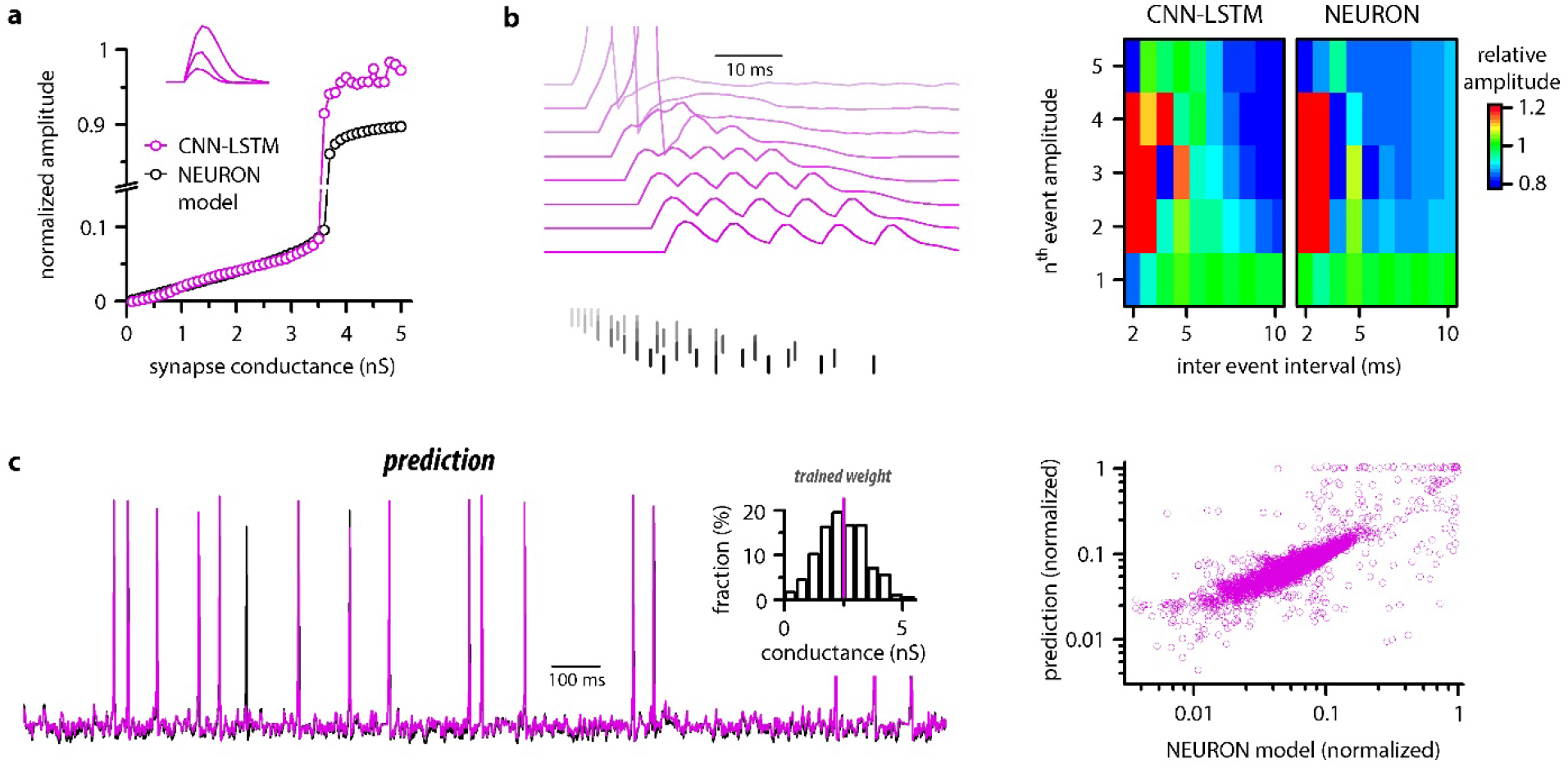
Ideal generalization of the CNN-LSTM trained on single input weight. **a**. CNN-LSTM models predict similar subthreshold event amplitudes and action potential threshold (break in y-axis) for increasing input weight, compared to NEURON models. **b**. CNN-LSTM models correctly represent temporal summation of synaptic events. Representative traces for different inter-event intervals (range: 2-10 ms, 1 ms increment) on the left, comparison of individual events in a stimulus train, relative to the amplitude of unitary events on the right. **c**. Single simulated active membrane potential trace in CNN-LSTM (purple) and NEURON (black) with variable synaptic input weights (left). The inset shows the distribution of synaptic weights used for testing generalization, with the original trained synaptic weight in purple. CNN-LSTM predicted membrane potential values plotted against NEURON model ground truth (right). Plotted values correspond to continuously predicted CNN-LSTM traces.

NEURON models can calculate and display several features of neuronal behavior in addition to membrane potential, including ionic current flux. To test how our CNN-LSTMs perform in predicting ionic current changes, we supplemented ANN inputs with sodium (I_Na_) and potassium currents (I_K_) and tasked the models to predict these values as well. The accuracy of the CNN-LSTM prediction for these ionic currents was similar to membrane potential predictions (Fig. 3, Pearson’s r = 0.999 and 0.99 for fitting, n = 5000, variance explained: 0.64 ± 0.21 and 0.55 ± 0.23, prediction correlation coefficient: 0.85 ± 0.08 and 0.81 ± 0.1, n = 5, for I_K_ and I_Na_, respectively) while the other ANNs again regressed to the mean. Together, these results demonstrate that the CNN-LSTM correctly learned several highly specialized aspects of neuronal behavior.

**Fig. 3.**
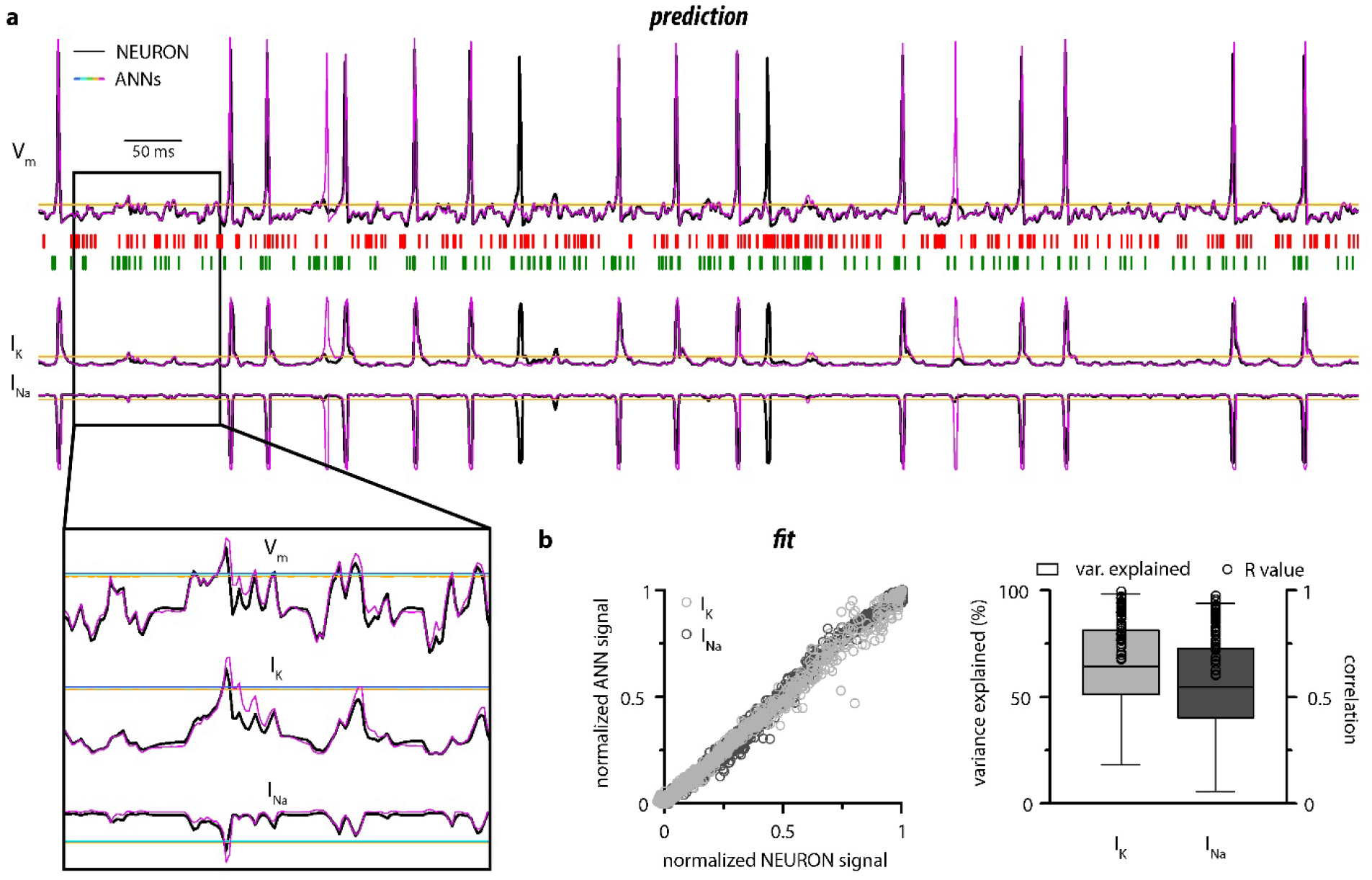
CNN-LSTM prediction of neuronal mechanisms beyond somatic membrane potential. **a**. Representative membrane potential (V_m_, top) and ionic current (I_K_ – potassium current, I_na_ – sodium current, bottom) dynamics prediction upon arriving excitatory (green, middle) and inhibitory (red, middle) events. Enlarged trace shows subthreshold voltage and current predictions. Color coding is same as for Figure 1. (black – NEURON model traces, magenta – CNN-LSMT, blue – linear model, teal – nonlinear model, green – deep neural net, orange – convolutional net). Notice the smooth vertical line corresponding to predictions by ANNs, with the exception of CNN-LSTM. On bottom left, magnified view illustrates the subthreshold correspondence of membrane potential and ionic current traces. **b**. CNN-LSTM models accurately predict ionic current dynamics. Normalized ANN predictions are plotted against normalized neuron signals for sodium (dark grey, left) and potassium currents (light grey). Variance of suprathreshold traces is largely explained by CNN-LSTM predictions (right, color coding is same as in panel B left). Correlation coefficients are superimposed in black.

### Predicting the activity of morphologically realistic neurons using ANNs

Neurons multiply their adaptive properties by segregating different conductances into separate subcellular compartments^59-66^. Thus, in addition to simplified input integrating point neurons, a substantial portion of neuronal models developed in recent decades intended to address subcellular signal processing via detailed multicompartmental biophysical cellular representations^15,42,65,67-69^. Therefore, our next aim was to examine how well ANNs describe multicompartmental information. To this end, a training dataset of synaptic inputs and corresponding somatic voltage responses was generated in NEURON from a morphologically and biophysically detailed *in-vivo* labeled neocortical layer 5 (L5) pyramidal cell (PC)^42^. We found that the trained CNN-LSTM performed in near-perfect accordance with the NEURON simulation (Fig. 4a, Pearson’s r = 0.999, n = 45000 ms). The continuous self-reliant prediction yielded lower yet adequate AP fidelity (Fig. 4g, 68.28 ± 18.97 % and 66.52 ± 25.37 % precision and recall, 0.439 ± 4.181 ms temporal shift for true positive spikes compared to ground truth, n = 205) compared to the point neuron, and the accuracy of subthreshold membrane potential fluctuations remained high (Pearson’s r = 0.83, n = 37).

**Fig. 4.**
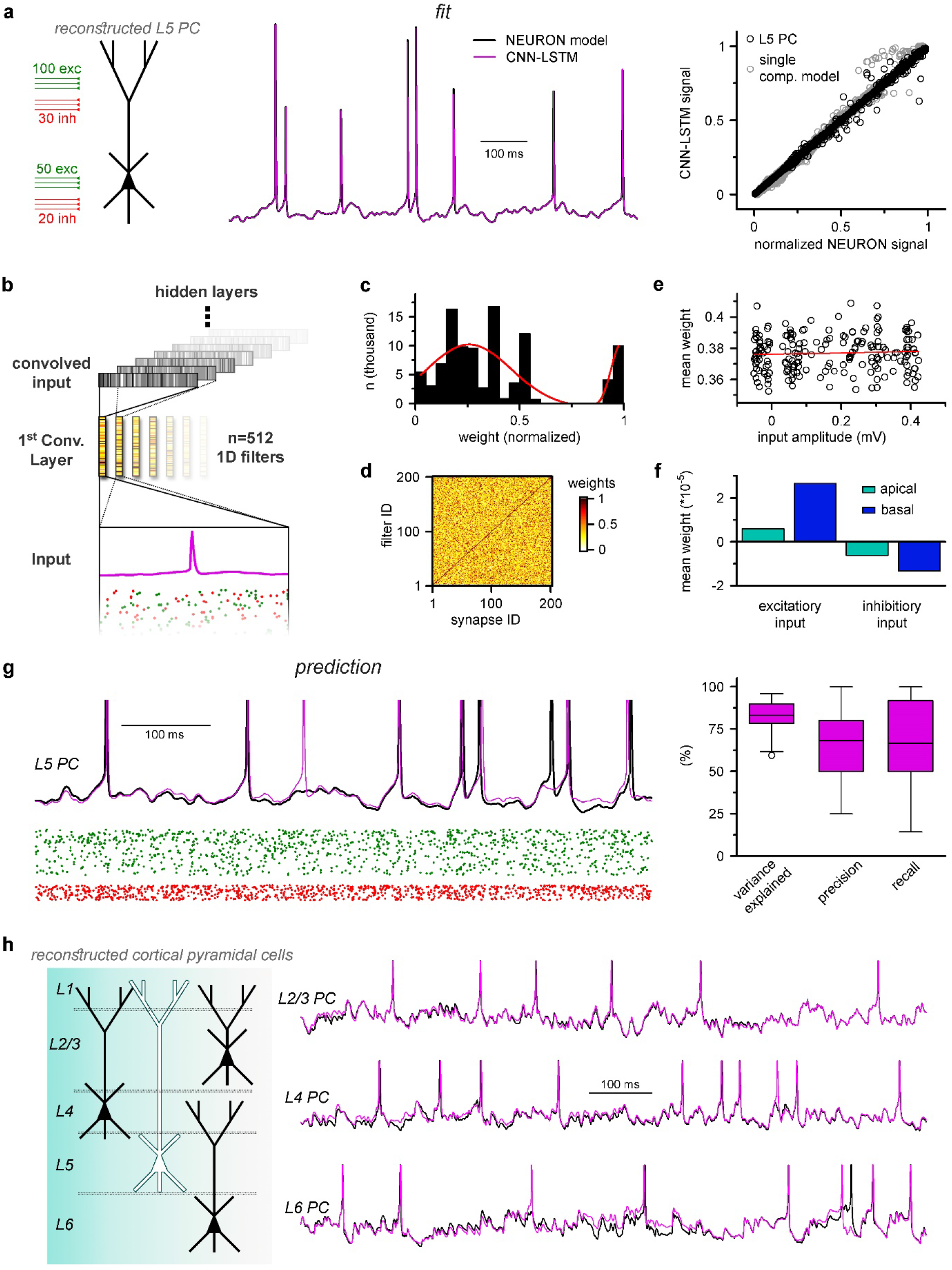
Multicompartmental simulation representation by CNN-LSTM. **a**. CNN-LSTM can accurately predict membrane potential of a multicompartmental neuron upon distributed synaptic stimulation. Representative figure depicts the placement of synaptic inputs (150 excitatory inputs: 100 inputs on apical, oblique and tuft dendrites and 50 inputs on the basal dendrite, randomly distributed, and 50 inhibitory inputs: 30 inputs on apical, oblique and tuft dendrites and 20 inputs on the basal dendrite, randomly distributed) of a reconstructed L5 PC (left). Point-by-point forecasting of L5 PC membrane potential by a CNN-LSTM superimposed on biophysically detailed NEURON simulation (left). CNN-LSTM prediction accuracy of multicompartmental membrane dynamics is comparable to single compartmental simulations (right, L5 PC in black, single compartmental simulation of Figure 1d in grey, n = 45000 and 50000 respectively). **b**. Convolutional filter information was gathered from the first convolutional layer (middle, color scale depicts the different weights of the filter), which directly processes the input (membrane potential in magenta, excitatory and inhibitory synapse onsets in green and red respectively), providing convolved inputs to upper layers (grey bars, showing the transformed 1D outputs). **c**. Distribution of filter weights from 512 convolutional units (n = 102400) with double Gaussian fit (red). **e**. Filter weight is independent of the somatic amplitude of the input (circles are averages from 512 filters, n = 200, linear fit in red). **d**. Each synapse has a dedicated convolutional unit, shown by plotting the filter weights of the 200 most specific units against 200 synapses. Notice the dark diagonal illustrating high filter weights. **f**. Excitatory and inhibitory synapse information is convolved by filters with opposing weights (n = 51200, 25600, 15360 and 10240 for apical excitatory, basal excitatory, apical inhibitory and basal inhibitory synapses respectively). **g**. Representative continuous prediction of L5 PC membrane dynamics by CNN-LSTM (magenta) compared to NEURON simulation (black) upon synaptic stimulation (left, excitatory input in green, inhibitory input in red). Spike timing is measured on subthreshold traces (right, n = 50 for variance explained, precision and recall). **h**. ANNs constrained on cortical layer 2/3 (top), layer 4 (middle) and layer 6 (bottom) pyramidal cells selected from the Allen Institute model database.

Establishing a proper multicompartmental representation of a neural system by relying solely on the somatic membrane potential is a nontrivial task due to complex signal processing mechanisms taking place in distal subcellular compartments^70-74^. This is especially true with respect to signals arising from more distal synapses^75-77^. To examine whether the CNN-LSTM took distal inputs into consideration or neglected these inputs in favor of more robust proximal ones, we inspected the weights of the first layer of the neural network architecture (Fig. 4b). This convolutional layer consists of 512 filters, which directly processes the input matrix (64 ms of 201 input vectors corresponding to the somatic membrane potential and vectorized timing information of 200 synapses). Despite the random initialization of these filters from a uniform distribution^78^, only a small fraction of optimized filter weights were selected for robust information representation (13.83% of all weights were larger than 0.85), while the majority of them were closer to zero (Fig. 4c) suggesting relevant feature selection. In order to demonstrate that this feature selection was not biased against distal inputs, the 512 convolutional filters were ranked by their selectivity for distinct synapses. We found that each synaptic input was assigned an independent selectivity filter (Fig. 4d). Next, we compared the mean weights of each synapse with the somatic amplitude of the elicited voltage response as a proxy for input distance from the soma (Fig. 4e). This comparison revealed a flat linear correspondence (Pearson’s r = 0.06), which combined with the filter specificity (Fig. 4d) confirmed that distal and proximal synaptic inputs carry equally relevant information for the CNN-LSTM.

Interestingly, when we compared the weights of excitatory and inhibitory inputs, we found that even at the first layer, the CNN-LSTM could determine that these inputs have opposing effects on subsequent membrane potential (5.91*10^−6^, 2.66*10^−5^, -6.22*10^−6^ and -1.34*10^−5^ mean weights for apical excitatory, basal excitatory, apical inhibitory and basal inhibitory synapses respectively, n = 51200, 25600, 15360 and 10240) even though these vectors only contain synaptic conductance information (comparable positive values for both excitatory and inhibitory synapses, Fig. 4f). Taken together, the feature selectivity and prediction accuracy confirm that the CNN-LSTM architecture is well suited for representing multicompartmental information.

We next aimed to establish a streamlined method to constrain ANNs on neuronal models from a publicly available (Allen Institute), well-curated database^79^ without developer involvement. Using this data pipeline, we constrained ANNs on other cortical pyramidal cell types; layer 2/3, layer 4, and layer 6 pyramidal cells (Fig. 4h). We found that the resulting ANNs again were fit adequately to ground truth NEURON simulations (60.9 ± 4.01 % and 63.17 ± 5.92 % precision and recall). Together, we have developed an ANN architecture appropriate for multicompartmental neuronal simulations of diverse cell types and laid the groundwork for a user-friendly methodology for their construction and integration into more complex circuit models.

### Ultra-rapid single neuron and network simulations using CNN-LSTM

One of the main reasons we chose a machine learning approach as a substitute for traditional modeling environments is the potential for markedly reduced simulation runtimes. Simulation environments such as NEURON rely on compartment-specific mathematical abstractions of active and passive biophysical mechanisms^54^, which results in high computational load in increasingly complex circuit models. In the case of moderately sized^80-84^ and full-scale networks^19-21,85^ this hinders the possibility of running these models on non-specialized computational resources. Although several attempts have been made to reduce the demanding computational load of neuronal simulations^32,33,86-90^, the most commonly used approach is parallelization, both at the level of single cells^91^ and network models^92,93^. ANNs offer a unique solution to this problem. Contrary to traditional modeling environments, graph based ANNs are designed explicitly for parallel information processing, meaning that run times should not increase linearly after additional cells are integrated into the simulated circuit (Fig. 5a). Thus, ANNs are well suited for scaling for large networks where an immense number of cells are simulated.

**Fig. 5.**
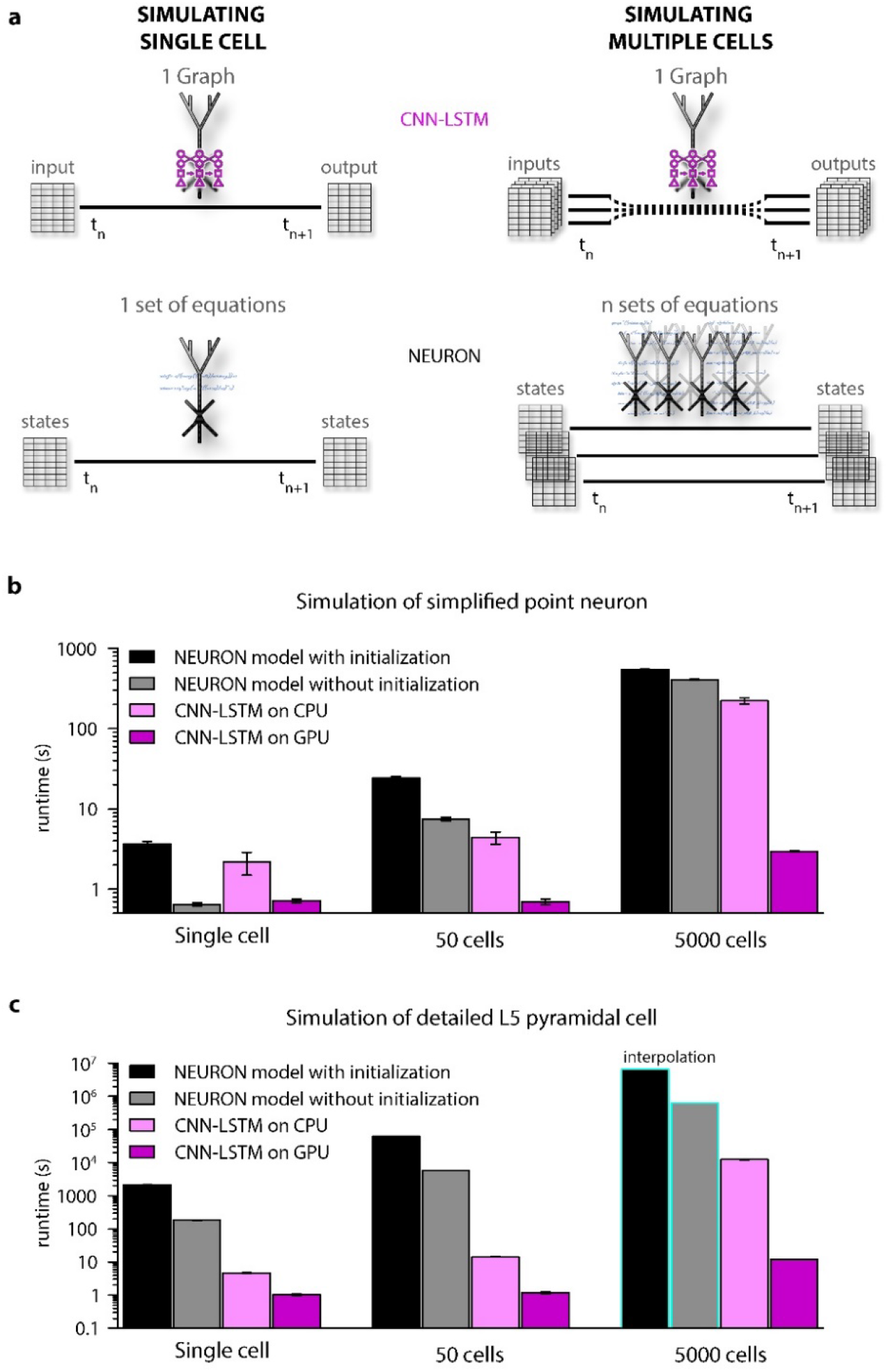
Orders of magnitude faster simulation times with CNN-LSTM. **a**. An illustration demonstrating that CNN-LSTMs (top, magenta) handle both single cell (left) and network (right) simulations with a single graph, while the set of equations to solve increases linearly for NEURON simulations (bottom, black). **b**. 100 ms simulation runtimes of a one-, 50- and 5000-point neurons using different resources. Bar graphs represent the average of five simulations. **C**. Same as in panel B, but for L5 PC simulations. Teal borders represent interpolated datapoints.

To verify the efficiency of our CNN-LSTM, we compared single cells with moderate-to large-scale network simulation runtimes against NEURON models used in Fig. 1 and Fig. 4. NEURON simulations were performed on a single central processing unit (CPU), as this is the preferred and most widely used method (although see^94,95^), while neural nets were tested on both CPU and graphical processing unit (GPU) because these calculations are optimized for GPUs. For point neurons, when the optional initialization step was omitted, single cell simulations ran significantly faster in NEURON than their CNN-LSTM counterparts (Fig. 5b, 3.68 ± 0.24 s, 0.65 ± 0.03 s, 2.19 ± 0.69 ms and 0.72 ± 0.04 s, 100 ms cellular activity by NEURON with initialization, NEURON without initialization, CNN-LSTM on CPU and CNN-LSTM on GPU respectively, n = 5). However, with increasing network size, the predicted optimal scaling of CNN-LSTM models resulted in faster runtimes compared to NEURON models for a 50-cell network (24.23 ± 1.12 s, 7.45 ± 0.37 s, 4.42 ± 0.77 s and 0.71 ± 0.05 s for simulating 100 ms network activity by NEURON with initialization, NEURON without initialization, CNN-LSTM on CPU and CNN-LSTM on GPU respectively, n = 5). These results show that in NEURON, runtimes increased by approximately 6.6-times, while CNN-LSTM runtimes on a GPU did not increase.

To demonstrate the practicality of ANNs for typical large-scale network simulations, we repeated these experiments with 5000 cells (representing the number of cells in a large-scale network belonging to the same cell type^21^). In these conditions, the NEURON simulation was ∼148 times slower than a single cell simulation. However, the large-scale CNN-LSTM simulation was only ∼4 times slower than that of a single cell (Fig. 5b, 546.85 ± 4.61 ms, 407.2 ± 9 ms, 222.15458 ± 19.02 ms, and 2.97 ± 0.02 ms for simulating 100 ms network activity by NEURON with initialization, NEURON without initialization, CNN-LSTM on CPU, and CNN-LSTM on GPU respectively.

We next compared runtime disparities for NEURON and CNN-LSTM simulations of detailed biophysical models. We found that the single cell simulation of the L5 PC model ran significantly slower than the CNN-LSTM abstraction (2.08*10^3^ ± 84.66 s, 185.5 ± 3.7 s, 4.73 ± 0.13 s and 1.02 ± 0.05 s for simulating 100 ms network activity by NEURON with initialization, NEURON without initialization, CNN-LSTM on CPU and CNN-LSTM on GPU respectively, n = 5). This runtime disparity was markedly amplified in network simulations (50 cell network: 6.3*10^4^ s, 5.8*10^3^ s, 14.3 ± 0.24 s and 1.19 ± 0.08 s, 5000 cell network: 6.53*10^6^ s, 6.28*10^5^ s, 901.15 s and 11.99 s for simulating 100 ms network activity by NEURON with initialization, NEURON without initialization, CNN-LSTM on CPU and CNN-LSTM on GPU respectively, n = 5), resulting in a four to five orders of magnitude faster runtime (depending on initialization) for the CNN-LSTM in case of large-scale network simulations. These results demonstrate our machine learning approach yields far superior runtimes compared to traditional simulating environments. Furthermore, this acceleration is comparable to that afforded by increased parallel CPU cores used for several network simulations^19-21^ introducing the possibility of running large or full-scale network simulations on what are now widely available computational resources.

### Efficient parameter space mapping using ANNs

Due to slow simulation runtimes, network simulations are typically carried out only a few times (but see^96^), hindering crucial network construction steps, such as parameter space optimization. Therefore, we investigated whether our ANN approach supports exploration of the parameter space in a pathophysiological system characterized by multidimensional circuit alterations, using Rett syndrome as a validation example. Rett is a neurodevelopmental disorder is caused by loss-of-function mutations in the X-linked methyl-CpG binding protein (MeCP2)^44^. Rett occurs in ∼1:10,000 births worldwide, resulting in intellectual disability, dysmorphisms, declining cortical and motor function, stereotypies, and frequent myoclonic seizures^97-103^. Although the underlying cellular and network mechanisms are largely unknown, changes in synaptic transmission^104-106^, neuronal morphological alterations^100,107^, and altered network connectivity^108^ have been reported in Rett syndrome.

We aimed to investigate the contribution of distinct alterations on cortical circuit activity in Rett syndrome, using a recurrent L5 PC network^82^ composed entirely of CNN-LSTM-L5-PCs (Fig. 6a). Simulations were run uninterrupted for 100 ms, when a brief (1 ms) perisomatic excitation was delivered to mimic thalamocortical input onto thick tufted pyramidal cells^109-111^. In control conditions, cells fired well-timed APs rapidly after the initial stimuli followed by an extended AP firing due to the circuit recurrent connectivity. (Fig. 6b,^112,113^). First, we compared the run time of the simulated layer 5 microcircuit of NEURON and CNN-LSTM models. We found that for a single simulation, CNN-LSTM models were more than 9300-times faster compared to NEURON models (Fig. 6c, 21.153 ± 0.26 s vs 54.69 h for CNN-LSTM and NEURON models respectively).

**Fig. 6.**
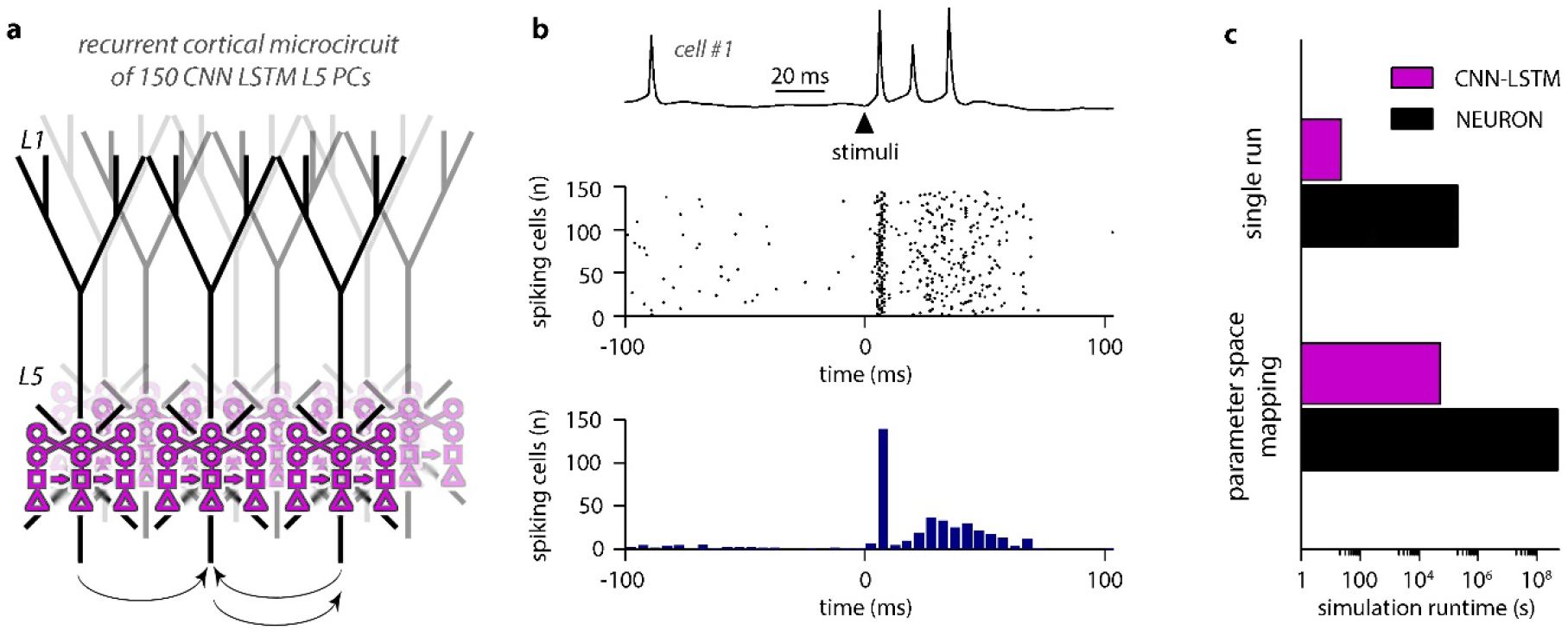
Efficient parameter-space mapping with CNN-LSTMs reveals a joint effect of recurrent connectivity and E/I balance on network stability and efficacy in Rett syndrome. **a**. 150 CNN-LSTM models of L5 PCs were simulated in a recurrent microcircuit. **b**. The experimental setup consisted of a stable baseline condition for 100 ms, a thalamocortical input at t = 100 ms, and network response, monitored for 150 ms. Example trace from the first simulated CNN-LSTM L5 PC on top, raster plot of 150 L5 PCs in the middle, number of firing cells with 5 ms binning for the same raster plot in the bottom. Time is aligned to the stimulus onset (t = 0, black arrowhead). **c**. Simulation runtime for single simulation (left, network of 150 cells simulated for 250 ms) and parameter space mapping (right, 150 cells simulated for 250 ms, 2500 times, for generating Figure 7 panel **b**).

### Rett cortical network alterations counteract circuit hyperexcitability

Cortical networks endowed with frequent recurrent connections between excitatory principal cells are prone to exhibit oscillatory behavior, which is often the mechanistic basis of pathophysiological network activities^114^. We quantified oscillatory activity^115-117^ and the immediate response to thalamocortical stimuli independently (Fig. 6c). By systematically changing excitatory quantal size^104^ and the ratio of recurrent L5 PC innervation, to mimic reduced recurrent connectivity and synaptic drive in Rett syndrome, we found that both alterations had considerable influence over network instability (Fig. 7b left panel; excitatory drive: 17.85 ± 61.61 vs 388.92 ± 170.03 pre-stimulus APs for excitatory drive scaled by 0.75 and 1.25, respectively, n = 100 each, p < 0.001; recurrent connectivity: 321.96 ± 200.42 vs 157.66 ± 192.5 pre-stimulus APs for 10% and 5.2% recurrent connectivity, similar to reported values for adult wild type and *Mecp2*-null mutant mice^108^, n = 50 each, p< 0.001) and response to stimuli (excitatory drive: 147.58 ± 17.2 vs 119.23 ± 18.1 APs upon stimulus for excitatory drive scaled by 0.75 and 1.25, respectively, n = 100 each, p = 2.3*10^−22^, t(198) = 11.03, *Two Sample t-Test*; recurrent connectivity: 134.76 ± 21.37 vs 112.74 ± 34.99 APs upon stimulus for 10% and 5.2% recurrent connectivity, n = 50 each, p = 2.54*10^−4^, t(98) = 3.8, *Two Sample t-Test*).

**Fig. 7.**
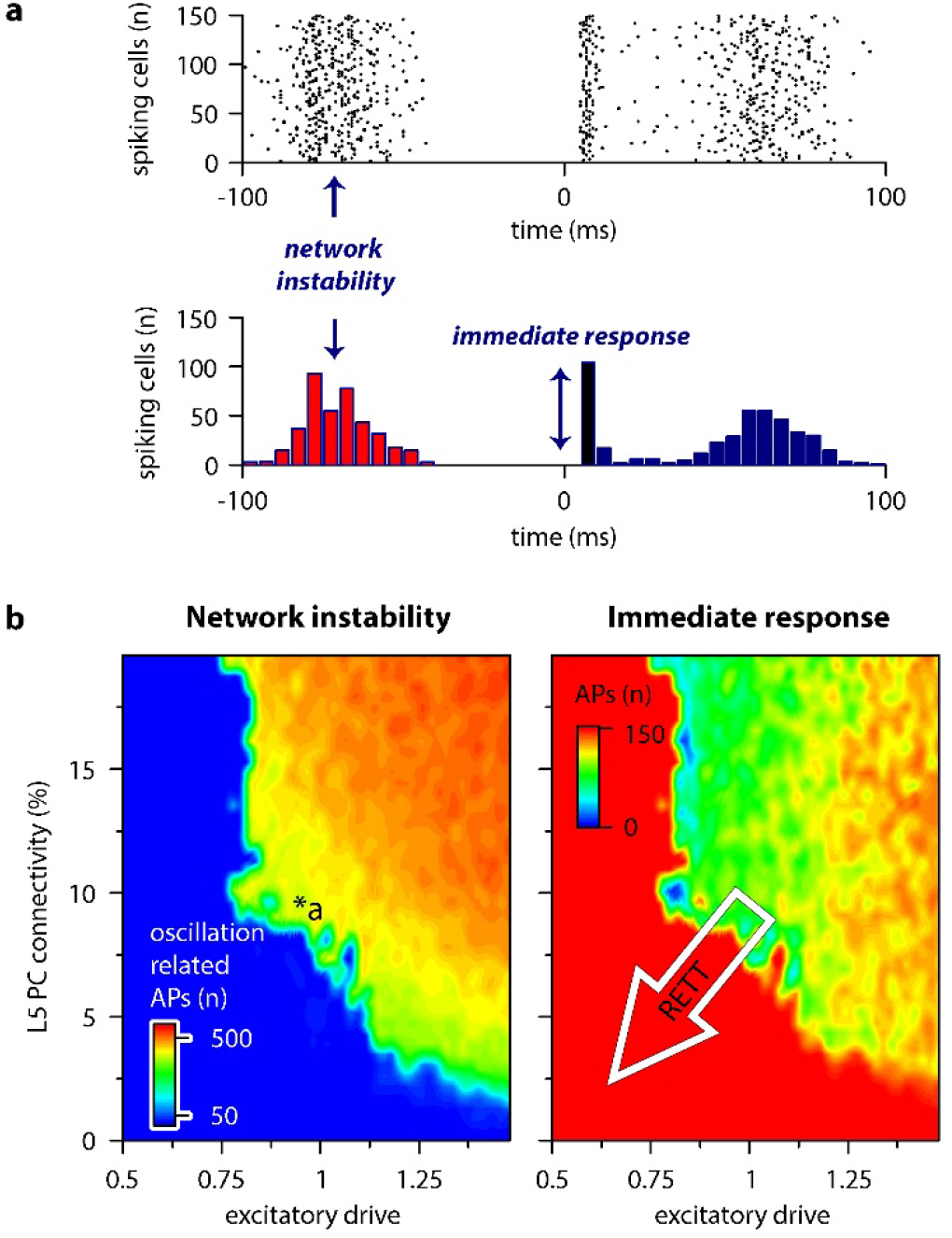
Recurrent connectivity and excitatory drive jointly define network stability in a reduced L5 cortical network. **a**. Two independent parameters were quantified: network instability (number of cells firing before the stimulus) and immediate response (number of cells firing within 10 ms of the stimulus onset). **b**. Network instability (left) and immediate response (right) as a function of altered L5 PC connectivity and excitatory drive. **\***a indicates network parameters used for generating panel a. The white arrow in the right panel denotes circuit alterations observed in Rett syndrome. Namely, 5% recurrent connectivity between L5 PCs instead of 10% in control conditions and reduced excitatory drive.

Contrary to disruption of the excitatory drive, when inhibitory quantal size^118^ was altered, we found that inhibition had only a negligible effect on network instability, as connectivity below 9% never resulted in oscillatory activity (Supplementary Fig. 2). Interestingly, we found no measurable relationship between the inhibitory quantal size and the network response to thalamocortical stimuli, which suggests that these metrics are robust for perturbations of the inhibitory circuitry in baseline conditions. These results suggest that lowered recurrent connectivity reduces network instability. Specifically, recurrent connectivity observed in young *Mecp2*-null mice (7.8 %^108^) yielded more stable microcircuits (54 % of networks were stable, n = 100), than wild-type conditions (34 % of networks were stable, n = 50). Recurrent connection probability of older animals (5.3%) further stabilized this network (64 % of networks were stable). Taken together, our model suggests that reduced recurrent connectivity between L5 PCs is not causal to seizure generation and abnormal network activity^103,115^, which are crucial symptoms of Rett syndrome at a young age. Rather, normal pyramidal cell activation is disrupted. This may correspond to the early stages of Rett syndrome where cortical dysfunction and regression are noted before the appearance of seizures^44^.

**Supplementary Fig. 2.**
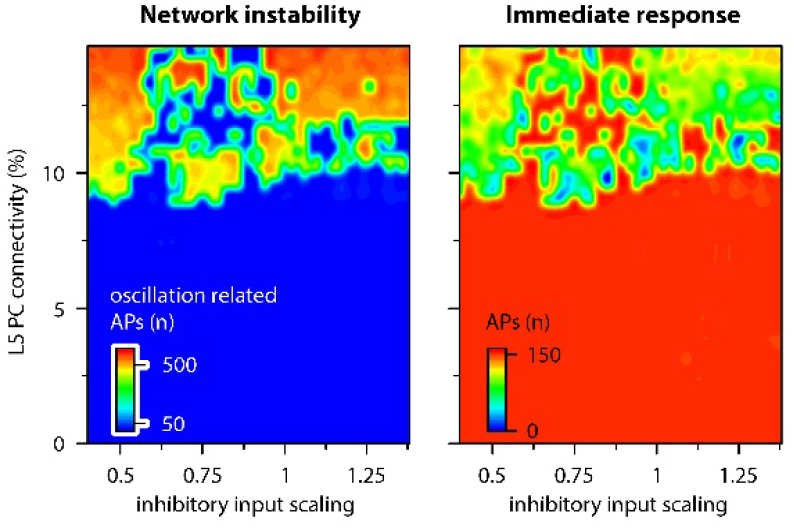
Microcircuit stability and efficacy is robust to changes in inhibitory drive. Network parameters were quantified as shown in Figure 7 panel **a**. Recurrent connectivity constrains network stability (9.14 ± 2.21 vs 320.78 ± 237.66 APs, n = 1740 vs 760 for below 9% connectivity and connectivity between 9-15 % respectively, p = 2.2*10^−219^, two sample t-test), while inhibitory inputs have a negligible effect (133.62 ± 29.32 vs 131.72 ± 32.32 APs upon thalamocortical stimulus for inhibitory input scaling of 1 and 0.5, respectively, n = 50 each, p = 0.76, t(98) = 0.31, Two Sample t-Test).

Together, we implemented multidimensional parameter space mapping in a cortical circuit exhibiting pathophysiological changes and identified the isolated outcome of distinct circuit alterations. Furthermore, our accelerated multicompartmental neural circuit model demonstrated that parameter space mapping is attainable by CNN-LSTM models, using commercially inexpensive computational resources.

## DISCUSSION

In this study, we present an ANN architecture (CNN-LSTM) capable of accurately capturing neuronal membrane dynamics. Most of the investigated ANN architectures predicted subthreshold voltage fluctuations of point-neurons, however only the CNN-LSTM could generate action potentials. This model could generalize well to novel input and could also predict various other features of neuronal cells, such as voltage-dependent ionic current dynamics. Furthermore, the CNN-LSTM accounted for most of the variance of subthreshold voltage fluctuations of biophysically realistic L5 PC models with excitatory and inhibitory synapses distributed along the entirety of the dendritic tree. The timing of the predicted action potentials closely matched the ground truth data. Importantly, we found that although single cell simulation runtimes are comparable, the CNN-LSTM has superior scaling for large network simulations. Specifically, in the case of mid-sized biophysically detailed networks (50 cells), ANNs were more than three orders of magnitude faster, while for large-scale networks (5000 cells) ANNs are predicted to be five orders of magnitude faster than traditional modeling systems. These accelerated simulation runtimes allowed us to quickly investigate a L5 PC network in distinct conditions, for example, to uncover network effects of altered connectivity and synaptic signaling observed in Rett syndrome. In our example Rett cortical circuit model, recurrent connectivity and excitatory drive jointly shape network stability and responses to sensory stimuli, showing the power of this approach in generating testable hypotheses for further empirical work. Together, the described model architecture provides a suitable alternative to traditional modeling environments with superior simulation speed for biophysically detailed cellular network simulations.

As our familiarity with neuronal circuits grows, so does the complexity of models tasked with describing their activity. Consequently, supercomputers are a regular occurrence in research articles that describe large-scale network dynamics built upon morphologically and biophysically detailed neuronal models^19-21^. Here we developed an alternative to these traditional models, which can accurately represent the full dynamic range of neuronal membrane voltages in multicompartmental cells, but with substantially accelerated simulation runtimes.

ANNs are ideal substitutes to traditional model systems for several reasons. First, ANNs do not require hard coding of the governing rules for neuronal signal processing. When an ANN is created, it serves as a blank canvas which can derive the main principles of input-output processing and neglect otherwise unimpactful processes^119-121^. The degree of simplification depends only on the ANN itself, not the developer, thereby reducing human errors. However, architecture construction and training dataset availability represent limiting steps in ANN development^122^. Fortunately, the latter issue is void as virtually infinite neuronal activity training datasets are now available for deep learning. On the other hand, as we have demonstrated, the former concern can significantly impede ANN construction. Although we have shown that markedly divergent ANN architectures can accurately depict subthreshold signal processing, we found only one that was suitable for both subthreshold and active membrane potential prediction. This does not mean that the presented architecture is the only possible ANN model for neural simulations, as the machine learning is a rapidly progressing field that frequently generates highly divergent ANN constructs^123^. The importance of the network architecture is further emphasized by our findings demonstrating that ANNs with comparable or even greater numbers of freely adjustable parameters could not handle suprathreshold information.

The prevailing CNN-LSTM architecture was proven suitable for depicting membrane potential and ionic current dynamics of both simplified and biophysically detailed neuronal models and generalized well for previously unobserved simulation conditions. Therefore, these results prove that ANNs are ideal substitutes for traditional model systems for representing various features of neuronal information processing in significantly accelerated simulations. Future architecture alterations should focus on the continued improvement of action potential timing and prediction.

Accelerated simulation runtimes are particularly advantageous for large-scale biological network simulations. These simulations can provide support for testing several network-related queries, such as pharmaceutical targeting and systemic interrogation of cellular level abnormalities in pathophysiological conditions^124-130^. However, widespread adaptation of large-scale network simulations is hindered by the computational demand of these models that can only be satisfied by the employment of supercomputer clusters^19-21^. Because these resources are expensive, they do not constitute a justifiable option for general practice. Importantly, we have shown that ANNs can provide a suitable alternative to traditional modeling systems, and that their simulation runtimes are superior due to the structure of the machine learning platform.

Traditional model systems linearly increase the number of equations to be solved for parallelly simulated cells, while ANNs can handle cells belonging to the same cell type on the same ANN graph^131^. In our network models (150 cells; Fig. 6), NEURON simulations yield 150-times higher number of linear equations every time step, while ANNs used the same graph for all simulated cells. For example, the Allen Institute recently published a computational model of the mouse V1 cortical area^21^, consisting of 17 different cell types (with the number of cells corresponding to these cell types ranging from hundreds to more than ten thousand), which means that a complete cortical area could be simulated using only 17 ANNs. We have demonstrated that even for small networks consisting of only 150 cells of the same type, ANNs are more than four orders of magnitude faster compared to model environments used in the aforementioned V1 simulations. As large-scale network simulations are typically run using several thousand CPU cores in parallel, the provided run time acceleration suggests that network simulations relying on ANNs could negate the need for supercomputers. Instead, these models can be run on commercially available computational resources such as personal computers with reasonable timeframes.

Another advantage of our approach is the utilization of GPU processing, which provides a substantially larger number of processing cores^132,133^. The runtime difference that can be attributed to the employed computational resource can be seen by comparing CNN-LSTM simulations on CPU and GPU (Fig. 5b, c), which yields below an order of magnitude faster simulations on GPU in the case of moderate size networks (50 cells) and approximately two orders of magnitude difference for large networks. However, our results demonstrated that cortical pyramidal cell networks simulations are at least four orders of magnitude faster than traditional modeling environments, which confirms that the disparity in the number of cores can only partially account for the observed runtime acceleration. Furthermore, the NEURON simulation environment does not benefit as much from GPU processing as ANN simulations^94,134^. These results confirm that the drastic runtime acceleration is the direct consequence of the parallelized graph-based ANN approach.

To demonstrate the superiority of ANNs in a biologically relevant network simulation, we mapped the effects of variable network parameters observed in Rett syndrome. Rett syndrome is a neurodevelopmental disorder leading to a loss of cognitive and motor functions, impaired social interactions, and seizures in young females due to loss of function mutations in the X-linked *MeCP2* gene^44^). Like many brain diseases, these behavioral alterations are likely due to changes in several different synaptic and circuit parameters. MeCP2-deficient mice exhibit multiple changes in synaptic communication, affecting both excitatory and inhibitory neurotransmission and circuit-level connectivity as well. Excitatory transmission is bidirectionally modulated by *MeCP2* knock-out^135,136^ and overexpression^137^, and long term synaptic plasticity is also impaired in MeCP2-deficient mice^138,139^. Inhibitory signaling is also altered in several different brain areas^104,105^. Importantly, synaptic transmission is not only affected at the level of quantal parameters, but at the number of synaptic connections, because MeCP2 directly regulates the number of glutamatergic synapses^136^. This regulation amounts to a 39% reduction of putative excitatory synapses in the hippocampus^136^, and a 50% reduction in recurrent excitatory connections between layer 5 pyramidal cells^108^. Here, using our ANN network model approach, we investigated how these diverse mechanisms might contribute to overall circuit pathology.

We found the ability of the network to respond to external stimuli is affected by both alterations in synaptic excitation and changes in the recurrent connectivity of layer 5 pyramidal cells, but that disruption of inhibitory transmission is not necessary to elicit network instability in Rett. These results are supported by previous findings showing that both constitutive^140^ and excitatory cell targeted^141^ *MeCP2* mutation leads to network seizure generation, as opposed to inhibitory cell targeted *MeCP2* mutation, which causes frequent hyperexcitability discharges but never seizures^118^. Furthermore, our results suggest that excitatory synaptic alterations in Rett affect both general network responses and network stability, which may serve as substrates to cognitive dysfunction and seizures, respectively. Taken together, these results reveal how cellular-synaptic mechanisms may relate to symptoms at the behavioral level. Importantly, investigation of the multidimensional parameter space was made possible by the significantly reduced simulation times of our ANN, as identical simulations with traditional modeling systems are proposed to be four orders of magnitude slower.

## METHODS

### Single-compartmental NEURON simulation

Passive and active membrane responses to synaptic inputs were simulated in NEURON (^54^, version 7.7, available at http://www.neuron.yale.edu/neuron/). Morphology (single compartment with length and diameter of 25 µm) and passive cellular parameters (*R*_*m*_: 1 kΩ/cm^2^, *C*_*m*_: 1 µF/cm^2^, *R*_*i*_: 35.4 Ω/cm) were the same for both cases and resting membrane potential was set to -70 mV. Additionally, the built-in mixed sodium, potassium and leak channel (^142^, based on the original Hodgkin-Huxley descriptions) was included in the active model (g_Na_: 0.12 pS/µm^2^, g_K_: 0.036 pS/µm^2^, g_leak_: 0.3 nS/µm^2^). Reversal potentials were set to 50 mV for sodium, -77 mV for potassium and -54.3 mV for leak conductance. Simulations were run with a custom steady state initialization procedure^143^ for 2 seconds, after which the temporal integration step size was set to 25 µs.

In order to simulate membrane responses to excitatory and inhibitory inputs, the built-in AlphaSynapse class of NEURON was used (excitatory synapse: *τ*: 2 ms, *g*_*pas*_: 2.5 nS, *E*_*rev*_: 0 mV, inhibitory synapse: *τ*: 1 ms, *g*_*pas*_: 8 nS, *E*_*rev*_: -90 mV). The number of synapses was determined by a pseudo-random uniform number generator (ratio of excitatory to inhibitory synapses: 8:3). Timing of individual synapses was also randomly picked from a uniform distribution. During the 10-second-long simulations the membrane potential, I_Na_ and I_K_ currents were recorded along with the input timings and weights and were subsequently saved to text files. Simulations were carried out in three different conditions. First, resting membrane potential was recorded without synaptic activity. Second, passive membrane potential was recorded. Third, active membrane potential responses were recorded with fixed synaptic weights.

### Multicompartmental NEURON simulation

Active multicompartmental simulations were carried out using an *in vivo*-labeled and fully reconstructed thick tufted cortical L5 PC^42^. The biophysical properties were unchanged, and a class representation was created for network simulations. Excitatory and inhibitory synapses were handled similarly to single-compartmental simulations. 100 excitatory (*τ*: 1 ms, *g*_*pas*_: 3.6 nS, *E*_*rev*_: 0 mV) and 30 inhibitory synapses (*τ*: 1 ms, *g*_*pas*_: 3 nS, *E*_*rev*_: -90 mV) were placed on the apical, oblique or tuft dendrites, 50 excitatory and 20 inhibitory synapses were placed on basal dendrites. The placement of the synapses was governed by two uniform pseudo-random number generators, which selected dendritic segments weighed by their respective lengths and the location along the segment (ratio: 2:1:1:1, for apical excitatory, apical inhibitory, basal excitatory and basal inhibitory synapses). Simulations were carried out with varied synaptic weights and a wide range of synapse numbers.

### ANN benchmarking

MTSF models are ideal candidates for modelling neuronal behavior in a stepwise manner, as they can be designed to receive information about past synaptic inputs and membrane potentials in order to predict subsequent voltage responses. These ANNs have recently been demonstrated to be superior to other algorithms in handling multivariate temporal data such as audio signals^144^, natural language^48^ and various other types of fluctuating time series datasets^145-147^. To validate the overall suitability of different ANN architectures tested in this paper for multivariate time series forecasting, we used a weather time series dataset recorded by the Max Planck Institute for Biogeochemistry. The dataset contains 14 different features, including humidity, temperature and atmospheric pressure collected every 10 minutes. The dataset was prepared by François Chollet for his book Deep Learning with Python (dataset preparation steps can be found on the Tensorflow website: https://www.tensorflow.org/tutorials/structured_data/time_series). All ANN architectures were implemented using the Keras deep-learning API (https://keras.io/) of the Tensorflow open-source library (version 2.3,^148^, https://www.tensorflow.org/), with Python 3.7.

The first architecture we implemented was a simple linear model consisting of three layers without activation functions; a Flatten layer, a Dense (fully connected) layer with 64 units and a Dense layer with 3 units. The second architecture was a linear model with added nonlinear processing. The model contained three layers identical to the linear model, but the second layer had a sigmoid activation function. The third model was a deep neural net with mixed linear and nonlinear layers. Similar to the first two models, this architecture had a Flatten layer and a Dense layer with 64 units as the first two layers, followed by nine Dense layers (units: 128, 256, 512, 1024, 1024, 512, 256, 128, 64, for the 9 Dense layers) with hyperbolic tangent (tanh) activation function and Dropout layers with 0.15 dropout rate. The last layer was the same Dense layer with 3 units as in case of the linear and nonlinear models. The fourth model was a modified version of the WaveNet architecture introduced in 2016^52^, implemented based on a previous publication^53^. The fifth and final architecture was a convolutional LSTM model^49^ which consists of three distinct functional layer segments. The lowest layers (close to the input layer) were three, one dimensional convolutional layers (Conv1D) with 128, 100 and 50 units, and causal padding for temporal data processing. The first and third layer had kernel size of one and the second had kernel size of 5. The first two layers had “rectified linear unit” (relu) activation functions, and the third layer had tanh activation, therefore the first two layers were initialized by He-uniform variance scaling initializers^78^, while the third layer was initialized by Glorot-uniform initialization (also known as Xavier uniform initialization)^149^. After flattening and repeating the output of this functional unit, a single Long-Short Term Memory layer (LSTM^50^) handled the arriving input, providing recurrent information processing. This layer had 128 units, tanh activation function, Glorot-uniform initialization and was tasked to return sequences instead of the last output. The final functional unit was composed of four Dense layers with 100 units, scaled exponential linear unit (selu) activations and accordingly, Lecun-uniform initializations^150^. The dropout rate between Dense layers was set to 0.15.

All benchmarked architectures were compiled and fitted with the same protocol. During compiling, the loss function was set to calculate mean squared error and the Adam algorithm^151^ was chosen as the optimizer. The maximum number of epochs was set to 20, however an early stopping protocol was defined to have a patience of 10, which was reached in all cases.

### Single compartmental simulation representation with ANNs

As neural nets favor processed data scaled between -1 and 1 or 0 and 1, we normalized the recorded membrane potentials and ionic currents. Due to the 1 Hz recording frequency, AP amplitudes were variable beyond physiologically plausible ranges, therefore peak amplitudes were standardized. The trainable time series data was consisting of 64 ms long input matrices with 3 or 5 columns (corresponding to membrane potential, excitatory input, inhibitory input and optionally I_Na_ and I_K_ current recordings) and target sequences were vectors with 1 or 3 elements (membrane potential and optional ionic currents). Training, testing and validation datasets were created by splitting time series samples 80-10-10%.

Benchmarking the five different ANN architectures proved that these models can handle time series data predicting with similar accuracy, however, in order to obtain the best results, several optimization steps of the hyperparameter space were undertaken. Unless it is stated otherwise, layer and optimization parameters were unchanged compared to benchmarking procedures. First, linear models were created without a Flatten layer, instead of which a TimeDistributed wrapper was applied to the first Dense layer. The same changes were employed in case of the nonlinear model and the deep neural net. The fourth, convolutional model had 12 Conv1D layers with 128 filters, kernel size of 2, causal padding tanh activation function and dilatation rates constantly increasing by 2^n^. We found that the best optimization algorithm for passive and active membrane potential prediction is the Adam optimizer accelerated with Nesterov momentum^152^, with gradient clipping set to 1. Although mean absolute error and mean absolute percentage error was sufficient for passive membrane potential prediction, the active version warranted the usage of mean squared error in order to put emphasis on APs. We found out that mechanistic inference of the full dynamic range of simulated neurons was a hard task for ANNs, therefore we sequentially trained these models in a specific order. First, we taught the resting membrane potential by supplying voltage recordings with only a few or no synaptic inputs. This step was also useful to learn the isolated shapes of certain inputs. Second, we supplied highly active subthreshold membrane traces to the models and finally we inputted suprathreshold membrane potential recordings. During the subsequent training steps, previous learning phases were mixed into the new training dataset in order to avoid the catastrophic forgetting of gradient based neural networks^153^.

### CCN-LSTM for multicompartmental simulation representation

Data preprocessing was done as described for single compartmental representations. Time series data for CNN-LSTM input was prepared as matrices having 201 rows for membrane potential and 200 synapse vectors, and 64 rows (64 ms long input). The CNN-LSTM architecture consisted of three Conv1d layers (512, 256 and 128 units), a Flatten layer, a RepeatVector, three LSTM layers (128 units each) and six Dense layers (128, 100, 100, 100, 100, 1 units). Activation functions and initializations were similar to the CNN-LSTM described above, except for the first Dense layer, which had relu activation function and He-uniform initialization. Additionally, Lasso regularization^154^ was applied to the first Conv1D layer. We found that the best optimizer for our purposes was a variant of the Adam optimizer based on the infinity norm, called Adamax^151^. Due to the non-normal distribution of the predicted membrane potentials, an inherent bias was present in our results, which was scaled by either an additional bias term, or a nonlinear function transformation.

Network construction was based on a previous publication^82^. Briefly, 150 L5 PC were simulated in a network with varying unidirectional connectivity, and bidirectional connectivity proportional to the unidirectional connectivity (P_bidirectional_ = 0.5 * P_unidirecional_). Reciprocal connections were 1.5-times stronger than unidirectional connections. The delay between presynaptic AP at the soma and the onset of the postsynaptic response was 1 ms measured from the AP peak. Each connection consisted of five proximal contacts. Compared to the original publication, we modified the parameters of the Tsodyks-Markram model^155^ used to govern synaptic transmission and plasticity. Based on a recent publication^96^, we set U (fraction on synaptic resources used by a single spike) to 0.38, D (time constant for recovery from depression) to 365.6 and F (time constant for recovery from facilitation) to 25.71. The simulation was run for 250 or 300 ms, which consisted of a pre-stimuli period (to observe the occurrence of structured activity patterns) for 100 ms, and a post-stimuli period (to quantify network amplification). The stimulus itself consisted of a strong excitatory input (can be translated to 50 nS) delivered to a proximal dendritic segment, calibrated to elicit APs from all 150 cells in a 10 ms long time window. Scaling of inhibitory inputs was carried out by changing inhibitory quantal size of background inputs, while scaling of excitatory drive affected quantal size of recurrent synaptic connections as well.

### Computational resources

We used several different commercially available and free-to-use computational resources to demonstrate the attainableness of large network simulations using neural networks. Single compartmental NEURON simulations were carried out on a CPU (Intel Core i7-5557U CPU @3.1 GHz). For multicompartmental NEURON simulations, we used the publicly available National Science Foundation funded High Performance Computing resource via the Neuroscience Gateway^156^. In contrast to NEURON models, ANN calculations are designed to run on GPUs rather than CPUs. Therefore, ANN models were run on the freely accessible Google Collaboratory GPUs (NVIDIA Tesla K80), Google Collaboratory tensor processing units (TPUs, designed for handling tensor calculations typically created by Tensorflow) and a single high-performance GPU (GeForce GTX 1080 Ti). For speed comparison we run these models on Google Collaboratory CPUs (Intel Xeon, not specified, @2.2 GHz) and the previously mentioned CPU as well. During NEURON and ANN simulations parallelization was only employed for Neuroscience Gateway simulations and ANN fitting.

### Statistics

Averages of multiple measurements are presented as mean ± SD. Data were statistically analyzed by ANOVA test using Origin software and custom written Python scripts. Normality of the data was analyzed with Shapiro-Wilks test. Explained variance was quantified as one minus the fitting error normalized by the variance of the signal^47^. For accuracy measurements APs were counted within 10 ms time window as true positive APs. Precision and recall were calculated based on the following equations:

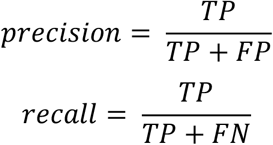

where FP in the false positive rate and FN is the false negative rate.

## Data and software availability

The code used for simulating single and multicompartmental NEURON models, ANN benchmarking, ANN representations and layer 5 microcircuit will be available upon publication.

## Acknowledgements

We would like to thank Annie Goettemoeller for her helpful comments on this manuscript. This work was supported by NIH grants R56AG072473 (MJMR), R21NS122011 (NPP), and K08NS105929 (NPP), and by CURE Epilepsy (NPP).

## Notes

### Competing Interest Statement

The authors have declared no competing interest.

